# Reaction cycle of operating pump protein studied with single-molecule spectroscopy

**DOI:** 10.1101/2021.05.24.445545

**Authors:** Saurabh Talele, John T. King

## Abstract

Biological machinery relies on nonequilibrium dynamics to maintain stable directional fluxes through complex reaction cycles. For such reaction cycles, the presence of microscopically irreversible conformational transitions of the protein, and the accompanying entropy production, is of central interest. In this work, we use multidimensional single-molecule fluorescence lifetime correlation spectroscopy to measure the forward and reverse conformational transitions of bacteriorhodopsin during trans-membrane H^+^ pumping. We quantify the flux, affinity, enthalpy and entropy production through portions of the reaction cycle as a function of temperature. We find that affinity of irreversible conformational transitions decreases with increasing temperature, resulting in diminishing flux and entropy production. We show that the temperature dependence of the transition affinity is well fit by the Gibbs-Helmholtz relation, allowing the ΔH_trans_ to be experimentally extracted.

## Introduction

Motor and pump proteins operate through non-thermal motions (nonequilibrium fluctuations)^[1–4]^ induced by the input of energy, for example, by ATP hydrolysis or photon absorption. For some pump proteins, the mechanism of action has been adequately described using the principle of microscopic reversibility,^[5]^ where equilibrium mechanical motions of the protein are rectified by an asymmetric potential that biases diffusion in one direction.^[6–7]^ In contrast, optically driven pump proteins are thought to operate through a ‘power-stroke’ mechanism, where a series of directional conformational transitions follow from a sudden structural change.^[3, 8]^ For this physical mechanism, which is inherently far-from-equilibrium, the validity of equilibrium notions of time-reversal symmetry and detailed balance, as well as the response to temperature, are yet to be established.

In this work, we study the reaction cycle of bacteriorhodopsin (bR), an optically-driven H^+^ pump found in Archaea (**Fig. 1a,b**),^[9]^ at the single-molecule level. This system was chosen as a model system as the intermediate species involved in the reaction cycle have been well characterized and bulk spectroscopy experiments have provided the timescale for elementary transitions of the cycle.^[10]^ However, ensemble experiments cannot resolve forward and reverse transitions required to quantify nonequilibrium thermodynamic properties.

**Figure 1.**
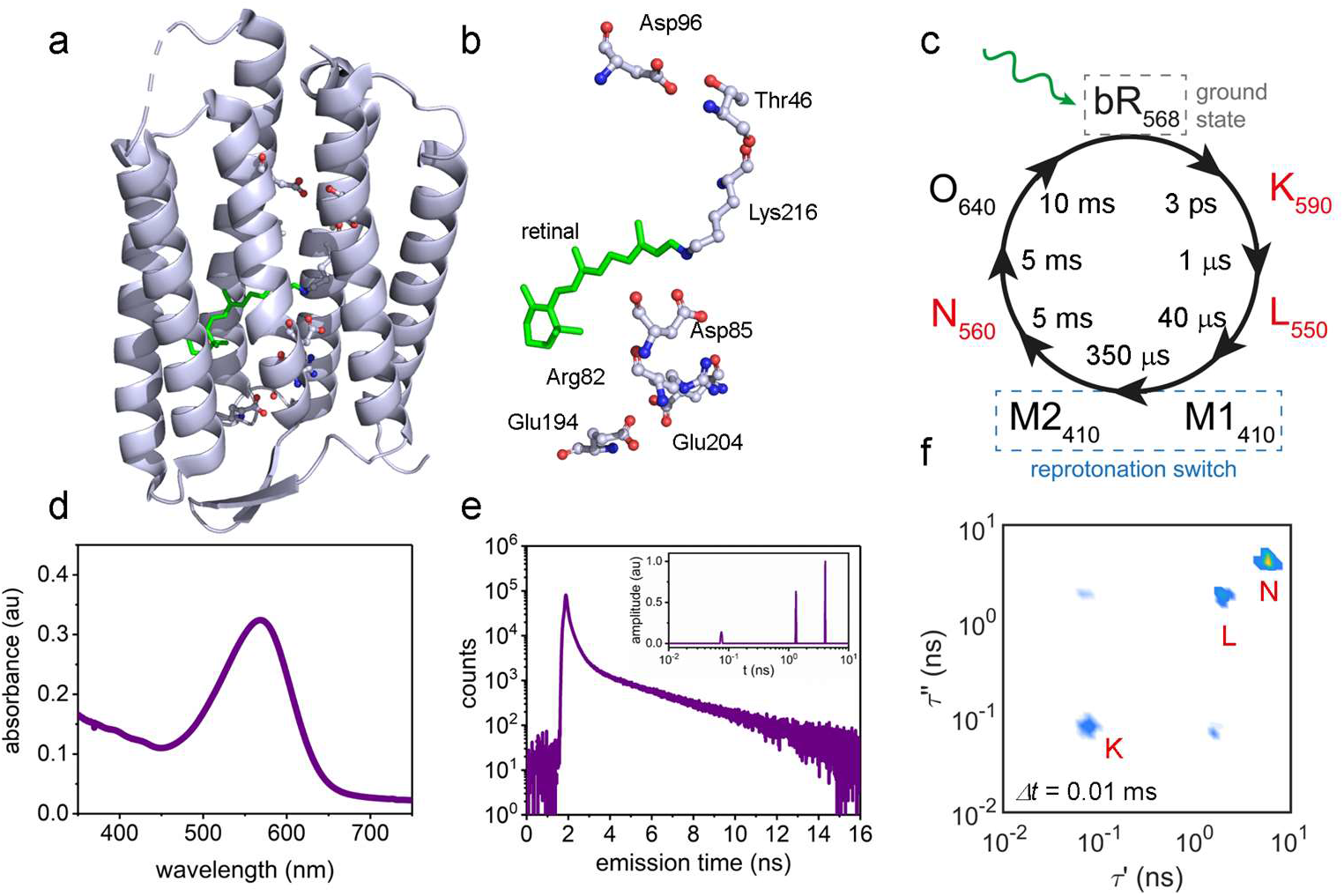
Single-molecule spectroscopy of bR. Crystal structure of ground state bR showing (**a**) the protein and (**b**) the retinal chromophore and the amino acids involved in the H^+^ transport chain of the reaction cycle (PDB: 1KBG). (**c**) Photo-induced reaction cycle of bR, which involves 6 intermediates in addition to the ground state (gs). The subscripts on the labels denote the absorption maximum of the retinal chromophore for the given conformation. (**d**) Absorption spectrum of retinal embedded in bR shows strong absorption at λ = 568 nm at pH = 6. (**e**) The fluorescence lifetime histogram measured from a single protein shows multi-component relaxation following 532 nm excitation. A 1D-ILT of the lifetime histogram reveals three relaxation times of τ_1_’ ~ 0.08 ns, τ_2_’ ~ 1.0 ns, and τ_3_’ ~ 5.0 ns (**inset**). (**f**) sm-2D-FLCS spectrum measured at a waiting time of *Δt* = 10 μs.

Photon absorption induces a *trans-cis* isomerization of the retinal chromophore. Steric repulsions between the isomerized chromophore and the protein scaffold initiates a multi-step reaction cycle involving multiple intermediate species (**Fig. 1c**).^[11–12]^ Key to our understanding of nonequilibrium reaction cycles are the roles of time-reversal symmetry and the corresponding entropy production.^[13]^ To characterize these properties on a single-molecule level, we leverage multidimensional single-molecule fluorescence lifetime correlation spectroscopy (sm-2D-FLCS),^[14–16]^ which utilizes multiple light-matter interactions separated by a controlled waiting time to monitor structural or chemical transitions of a molecule. The details of the technique are published elsewhere.^[17]^ Briefly, sm-2D-FLCS takes advantage of the differences in fluorescence lifetimes as a probe for resolving multiple intermediates during the catalytic cycle. The experiment is carried out on a confocal microscope equipped with TCSPC (Time Correlated Single Photon Counting) setup. Time resolved fluorescence photon streams are collected from single molecules and a 2D emission delay histogram is constructed by correlating the observed fluorescence emission delays in the photon stream at a known waiting time (*Δt*). A 2D Inverse Laplace transform (2D ILT) then yields a 2D fluorescence lifetime correlation spectra where species are represented by peaks corresponding to their fluorescence lifetimes. The exchange dynamics are observed through time-correlation functions (tcf) for forward and reverse transitions, which appear in opposite quadrants of the 2D spectrum^[18]^ and therefore allow the forward and reverse conformational transitions to be measured independently. The violation of time-reversal symmetry within the reaction cycle is observed from the reciprocal tcf,^[19]^ (where, reciprocal refers to the pair of forward and reverse tcf), which are equivalent for reversible transitions (obey time-reversal symmetry) and are not equivalent for irreversible transitions (violate time-reversal symmetry). The extent to which time-reversal symmetry is violated is a measure of entropy production rate in the cycle.^[13]^ Using this approach, we are able to experimentally characterize the nonequilibrium thermodynamics and kinetics of portions of the bR reaction cycle and quantify the stability of the cycle when subject to kinetic perturbation.

## Results and Discussion

We first characterize the directional flux through portions of the cycle by monitoring the transition kinetics between several intermediates. The endogenous retinal chromophore serves as the photo-trigger for the reaction cycle as well as a probe of the intermediates during the cycle. Limitations in the experiment prevent transitions between each intermediate to be directly measured. Instead, portions of the reaction cycle are characterized through select intermediates to which our experiments are sensitive. The cycle of bR involves six known intermediate structures that exist for timescales ranging between μs to ms (K, L, M1, M2, N, O, **Fig. 1c**).^[11–12]^ In the ground state structure, the retinal chromophore has an absorption maximum at λ_max_ = 568 nm (**Fig. 1d**). Fluorescence emission from a single monomeric bR protein, confirmed by diffraction limited emission spots observed in confocal microscopy and single-step photo-physics (**Fig. S1a-c**), shows multi-component relaxation following a 532 nm excitation (**Fig. 1e, Fig. S1d**). An inverse Laplace transform (ILT)^[17]^ of the cumulative photon histogram measured from more than 100 single protein molecules (**Fig. 1e**) gives a 1D fluorescence lifetime spectrum that contains three distinct relaxation peaks (**Fig. 1e, inset**). We do not anticipate signal from the M1, M2, and O intermediates due to low absorption at the 532 nm excitation.^[11–12]^ Furthermore, fluorescence from the ground state structure is unlikely due to the ultrafast isomerization reaction that occurs in the excited state.^[20]^ Therefore, the signals arise from the K intermediate, the L intermediate, and the N intermediate.

The 2D-FLCS spectrum generated from a 2D-ILT of the photon histogram^[17]^ calculated at a waiting time of Δt = 10 μs shows three distinct diagonal peaks (**Fig. 1f**). The cross-peaks between the signals at τ’ ~ 0.08 ns and τ’ ~ 1.0 ns indicate rapid exchange between the two states represented by these signals. However, no cross-peaks are observed between the diagonal signal at τ’ ~ 5.0 ns and the other two states. This is consistent with ensemble experiments which have measured the formation time of the N intermediate to be on the ms timescale.^[11–12]^ Furthermore, the 1D relaxation spectrum (**Fig. S3**) shows the peaks at τ’ ~ 0.08 ns, τ’ ~ 1.0 ns are in a 1:3 ratio, consistent with the populations of the K and L intermediates.^[21]^ From these observations we can assign the diagonal peaks at τ’ ~ 0.08 ns, τ’ ~ 1.0 ns, and τ’ ~ 5.0 ns to the K state, the L intermediate, and the N intermediates, respectively (**Fig. 1c, 1f**).

To characterize the kinetics of exchange between different intermediates of bR reaction cycle, we measure kinetic traces for variable waiting times *Δt* ranging from 0.01 to 200 ms (**Figure 2a**) at temperature of 291 K. The time-dependent amplitudes of the spectrum are given by the tcfs.^[14–15]^

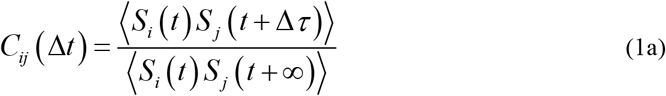

and

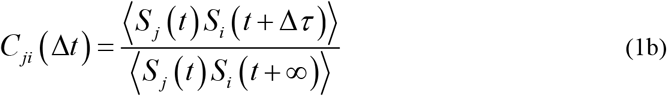

where *S_i_* and *S_j_* are the instantaneous probabilities of observing the system in states *i* and *j*, at given times respectively, measured in our experiment through unique fluorescence lifetimes *τ_i_* and *τ_j_* of each state. The equivalency of **Eq. 1a** and **Eq. 1b** under the condition of microscopic reversibility is given by detailed balance and time-reversal symmetry.^[19]^ The characteristic decay rate of the tcf for the *i* to *j* transition, *r_ij_*, is related to the transition probability by^[19]^

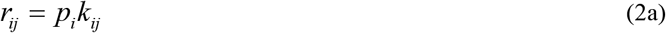

and

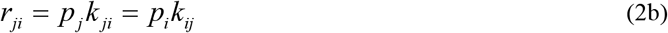

where *p_i_* is the average occupancy of state *i*, and *k_ij_* is the probability of observing a transition from state *i* to *j* within time *Δt*. Experimentally measured decay rate, *r_ij_*, and state occupancy *p_i_* allows for the transition probability *k_ij_* to be determined and an elementary rate law to be defined. Under conditions of microscopic reversibility, the forward and reverse transition probabilities are equivalent, reflecting detailed balance and time-reversal symmetry of the reaction process. The 2D spectra for a reversible process are therefore symmetric along the diagonal, as the reciprocal crosscorrelation functions are identical (defined by Eq. 1a and 1b). In contrast, for microscopically irreversible transitions, time-reversal symmetry of the reaction process is violated and the equivalency of cross-correlation functions Eq. 2a and Eq. 2b is broken. The resulting 2D spectra are asymmetric along the diagonal, indicating the presence of nonequilibrium fluxes through a given transition^[19, 22]^

**Figure 2.**
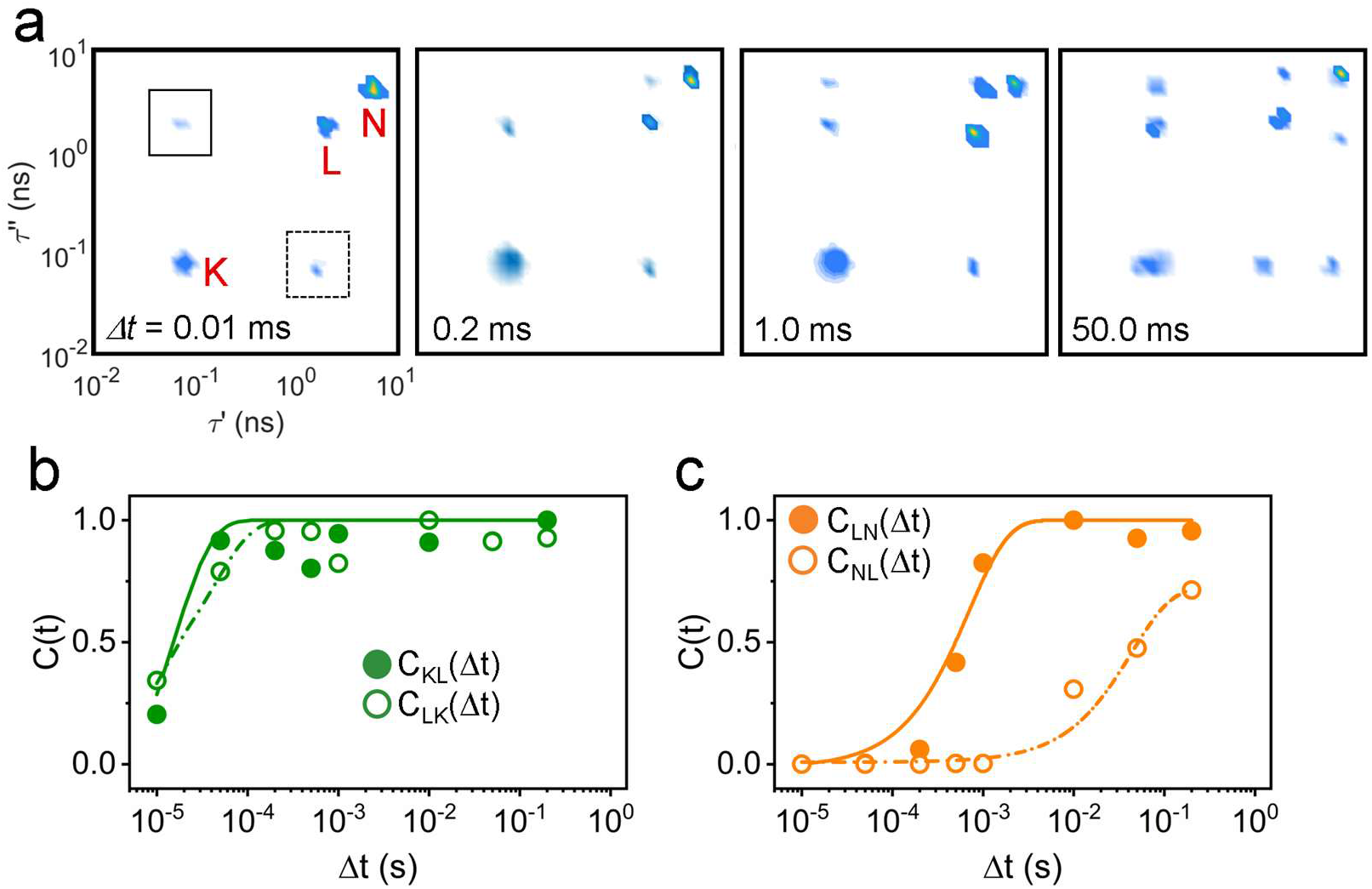
Microscopic irreversibility of bR catalytic cycle. (**a**) sm-2D-FLCS spectra of bR shown for waiting times ranging from 0.01 to 50 ms. Exchange kinetics at 291 K for waiting times *Δt* = 0.01 – 200 ms between (**b**) K intermediate to L intermediate and (**c**) L intermediate to N intermediate. Forward transitions, represented by solid symbols and solid lines, are measured from the upper quadrant of the 2D spectra. Reverse transitions, represented by open symbols and dashed lines, are measured from the lower quadrant of the 2D spectra. The forward (*k_ij_*) and reverse (*k_ji_*) transition rates are measured by fitting the cross-correlation functions to exponential growth functions (solid and dashed lines). The experiments were performed at T = 291 K.

The cross-peak dynamics between the K intermediate and the L intermediate show rapid forward (**Fig. 2b**, solid symbols) and reverse reactions (**Fig. 2b**, open symbols), with the exchange rates being of the same order of magnitude (*k_KL_* = 6.9×10^4^ s^−1^ and *k_LK_* = 6.6×10^4^ s^−1^). The near equivalency of the exchange rates suggests rapid equilibration of the transition. At *Δt* = 0.20 ms, a cross-peak emerges between the L and N intermediates that is asymmetric (no crosspeak for the reverse transition) (**Fig. 2a**), indicating a microscopically irreversible step in the reaction cycle. The forward transition occurs on a timescale of ~ 0.50 ms (*k_LN_* = 2.0×10^3^ s^−1^), while the reverse transition occurs on a timescale over 200 ms (*k_NL_* = 5.0 s^−1^) (**Fig. 2c**).

Chemically, the L-N transition corresponds to the reprotonation of the Asp96 residue on the cytoplasmic side of the protein (M intermediate) and formation of a Grotthuss-like H^+^ wire between the Asp96 residue and the Schiff base of retinal.^[23–24]^ Proton uptake and reprotonation of the retinal Schiff base (formation of N intermediate) from the cytoplasmic side is thought to be the switch step of the photo-cycle.^[25–26]^ The properties of the measured tcf demonstrate that the reprotonation of Asp96 is microscopically irreversible and has non-zero entropy production associated with the transition. The presence of microscopically irreversible transitions implies that nonequilibrium transitions are inherent to the protein’s reaction cycle and are required to maintain stable fluxes through portions of the cycle.

Next, we analyze the nonequilibrium thermodynamic properties of the bR reaction cycle as a function of temperature. The temperature-dependent occupancy of the K, L, and N states are shown in **Figure S3**. The *k_ij_* and *k_ji_* for an elementary transition are given by Arrhenius expressions,

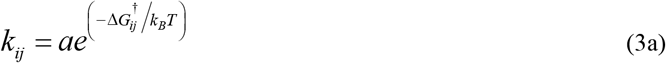

and

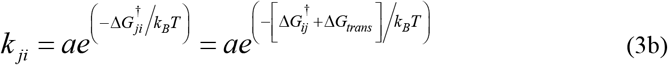

where *a* is a pre-exponential factor, *ΔG_ij_^†^* and *ΔG_ji_^†^* are the Gibbs free energy of activation for the forward and reverse transitions, respectively, and *ΔG_trans_* is the Gibbs free energy of the transition. Assuming the entropic contribution is temperature independent over the range studied, the temperature dependence of *k_ij_* and *k_ji_* for a given transition in the cycle will be identical on the condition that *ΔH_trans_* = 0 and different when *ΔH_trans_* ≠ 0. Arrhenius plots for the K-L transitions and the L-N transitions are shown in **Fig. 3a, b**. The *k_KL_* and *k_LK_* for the K-L transitions show identical temperature dependence (**Fig. 3a**), indicating that the *ΔH_trans_* = 0 for the transition. In contrast, the *k_LN_* and *k_NL_* for the L-N transitions (**Fig. 3b**) are significantly different at low temperature, but converge at high temperature, suggesting that *ΔH_trans_* ≠ 0 and the irreversibility of the transition needed to maintain a net forward flux through the cycle diminishes with increasing temperature.

**Figure 3.**
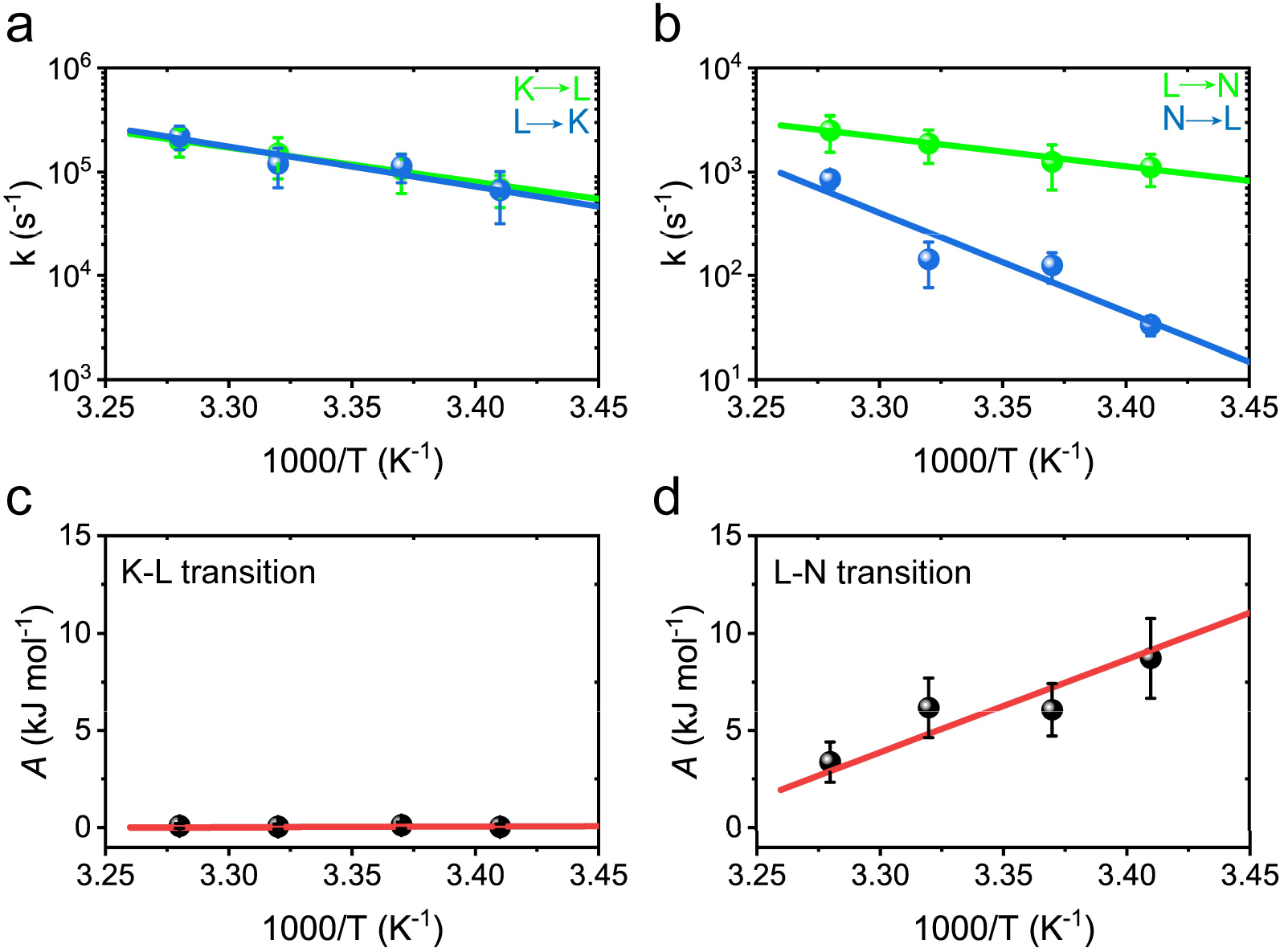
Thermal stability of the bR reaction cycle. Arrhenius plots for the (**a**) K-L and (**b**) L-N transition rates. The *k_ij_* and *k_ji_* values are shown in green and blue, respectively. (**c**) The affinity *A* of the K-L transition plotted vs T^−1^. As the transition is microscopically reversible, the *A* value is ~ 0 kJ mol^−1^ at all temperatures measured. (**d**) The *A* of the L-N transition plotted vs T^−1^. The microscopically irreversible transition has an *A* value of ~ 10 kJ mol^−1^ at 291 K that decreases linearly towards 0 kJ mol^−1^ with increasing temperature. The temperature dependencies of *A* are fit to the Gibbs-Helmholtz equation (red line, **Eq. 6**).

The affinity *A* of a nonequilibrium process quantifies the thermodynamic driving force. de Donder demonstrated that *A* is related to *ΔG^o^* for a chemical reaction.^[27]^

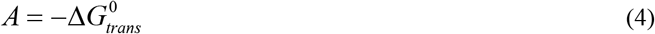

Using **Eq. 2–3**, we can rewrite *A* for a given transition in terms of *k_ij_* and *k_ji_*, and the state occupancies,

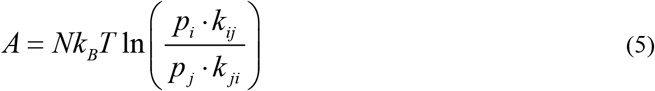

where the *ΔG^0^* values are given by *ΔG* – *Nk_B_Tlog(Q*), with *N* as Avagadro’s number and *Q* as the reaction quotient given by the ratio of state occupancies. Plots of *A* for the K-L and L-N transitions are shown in **Fig. 3c, d**. The *A* for the K-L transition, and by extension the *ΔG_trans_*, is 0 kJ mol^−1^ at all temperatures (**Fig. 3c**). In contrast, *A* for the L-N transition is ~ 10 kJ mol^−1^ at 291 K and decreases rapidly towards 0 kJ mol^−1^ with increasing temperature (**Fig. 3d**). According to **Eq. 4**, the temperature dependence of *A* is given by the Gibbs-Helmholtz equation.

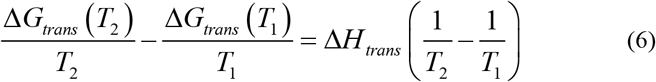

We find that the experimental data for *A* as a function of *T* is adequately fit with **Eq. 6**, yielding *ΔH_trans_* ~ 0 kJ mol^−1^ and *ΔH_trans_* ~ −30 kJ mol^−1^ for the K-L and L-N transitions, respectively (**Figure 3c, d**).

The thermodynamics of a nonequilibrium process can be quantified by the rate of entropy production rate, *σ*, given by ^[28]^,

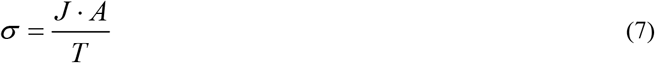

where *J* is the net flux defined for an elementary transition as the difference in forward and reverse reaction rates, *r_ij_* – *r_ji_*. Therefore, experimental measurements of the forward and reverse reaction rates (*r_ij_* and *r_ji_* and rate constants (*k_ij_* and *k_ji_*) are sufficient to determine *σ* of a nonequilibrium chemical process. For K-L and L-N transitions, *σ* is shown as a function of temperature (**Fig. 4a**). As expected, the K-L transition shows negligible *σ_KL_*. In contrast, the L-N transition shows significant *σ_LN_*, expected for an irreversible transition, that decreases exponentially with increasing temperature.

**Figure 4.**
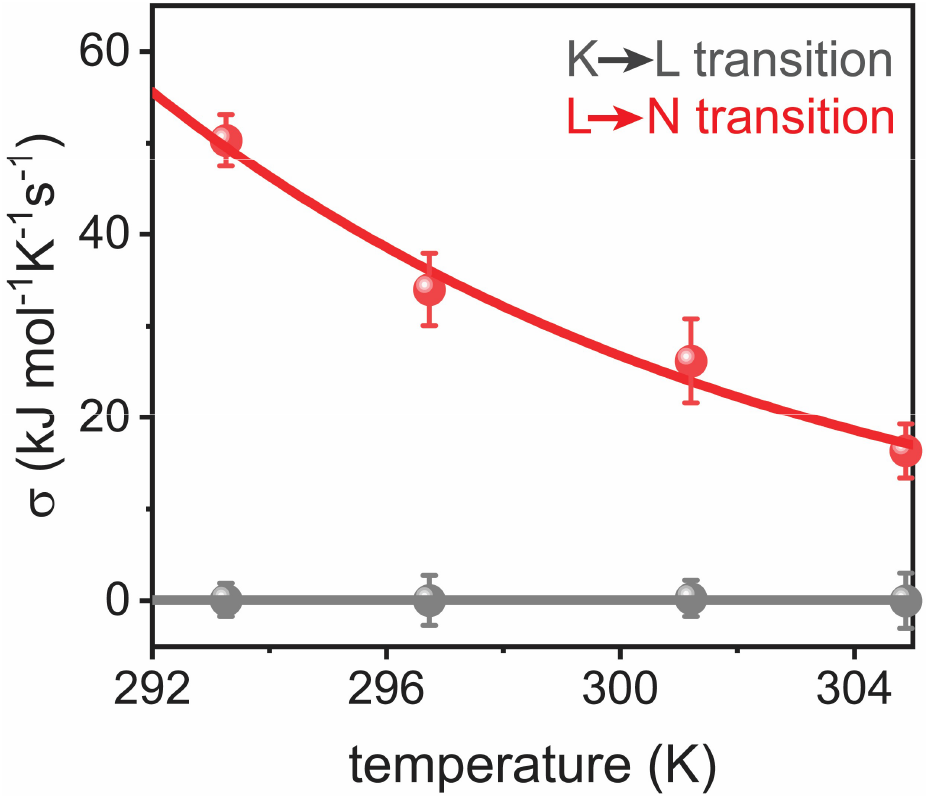
Entropy production rate and variance of flux. (**a**) Entropy production for the K-L and L-N transitions computed from **Eq. 7**. The entropy production rate *σ* associated with the reversible K-L transition is 0 kJ mol^−1^K^−1^s^−1^ at all temperatures measured, while the irreversible L-N transition has significant entropy production rate (~ 50 kJ mol^−1^K^−1^s^−1^) that decreases exponentially with increasing temperature.

## Conclusions

The mechanism by which biological machinery overcomes randomizing thermal forces to achieve directional action are not comprehensively understood, with key questions regarding microscopic reversibility and the applicability of equilibrium thermodynamics still remaining. The results presented here demonstrate that microscopic irreversibility is the fundamental operating principle of directional H^+^ transport by bacteriorhodopsin. Therefore, equilibrium notions of detailed balance and time-reversal symmetry often invoked for chemically driven pumps are not applicable to the function of optically driven pump proteins. Similarly, the thermal destabilization of the reaction cycle of bacteriorhodopsin is likely unique to pumps that operate through a power-stroke mechanism in which the nonequilibrium nature of the cycle is inherent to select conformational transitions of the protein.

## ACKNOWLEDGEMENTS

We thank F. Amblard and A. Shakya for productive discussions. We also thank the Korean taxpayers for supporting this work through the Korean Institute for Basic Science, Project Code IBS-R020-D1.

## ToC Information

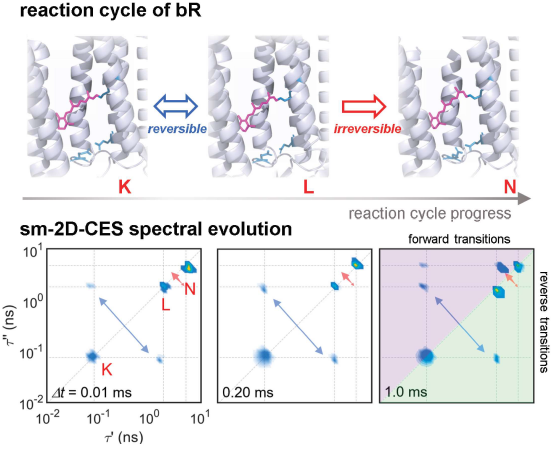

Pump proteins rely on the input, transmission, and dissipation of energy to carry out directional transport of ions across a membrane. In this work, we use multidimensional single–molecule spectroscopy to quantify the nonequilibrium thermodynamics for portions of the reaction cycle of bacteriorhodopsin. We demonstrate the role of microscopic irreversibility in the mechanism of action for optically driven pumps.

## REFERENCES

[1] R. D. Vale, R. A. Milligan, Science 2000, 288, 88–95.

[2] S. Leibler, D. A. Huse, J. Cell Biol. 1993, 121, 1357–1368.

[3] W. Hwang, M. Karplus, Proc. Natl. Acad. Sci. U.S.A. 2019, 116, 19777–19785.

[4] Y. Taniguchi, M. Nishiyama, Y. Ishii, T. Yanagida, Nat. Chem. Biol. 2005, 1, 342–347.

[5] R. D. Astumian, Nat. Nanotechnol. 2012, 7, 684–688.

[6] R. D. Astumian, Science 1997, 276, 917–922.

[7] F. Julicher, A. Ajdari, J. Prost, Rev. Mod. Phys. 1997, 69, 1269–1281.

[8] R. D. Astumian, Faraday Discuss. 2016, 195, 583–597.

[9] H. Luecke, B. Schobert, H. T. Richter, J. P. Cartailler, J. K. Lanyi, J. Mol. Biol. 1999, 291, 899–911.

[10] R. Neutze, E. Pebay-Peyroula, K. Edman, A. Royant, J. Navarro, E. M. Landau, Biochim. Biophys. Acta Biomembr. 2002, 1565, 144–167.

[11] J. K. Lanyi, Annu. Rev. Physiol. 2004, 66, 665–688.

[12] O. P. Ernst, D. T. Lodowski, M. Elstner, P. Hegemann, L. S. Brown, H. Kandori, Chem. Rev. 2014, 114, 126–163.

[13] C. Maes, K. Netocny, J. Stat. Phys. 2003, 110, 269–310.

[14] K. Ishii, T. Tahara, J. Phys. Chem. B 2013, 117, 11414–11422.

[15] K. Ishii, T. Tahara, J. Phys. Chem. B 2013, 117, 11423–11432.

[16] T. Otosu, K. Ishii, T. Tahara, Nat. Commun. 2015, 6, 1–9.

[17] S. Talele, J. T. King, Biophys. J. 2021, 120, 4590–4599.

[18] W. P. Aue, E. Bartholdi, R. R. Ernst, J. Chem. Phys. 1976, 64, 2229–2246.

[19] I. Z. Steinberg, Biophys. J. 1986, 50, 171–179.

[20] P. Nogly, T. Weinert, D. James, S. Carbajo, D. Ozerov, A. Furrer, D. Gashi, V. Borin, P. Skopintsev, K. Jaeger, K. Nass, P. Bath, R. Bosman, J. Koglin, M. Seaberg, T. Lane, D. Kekilli, S. Brunle, T. Tanaka, W. T. Wu, C. Milne, T. White, A. Barty, U. Weierstall, V. Panneels, E. Nango, S. Iwata, M. Hunter, I. Schapiro, G. Schertler, R. Neutze, J. Standfuss, Science 2018, 361.

[21] C. Wickstrand, P. Nogly, E. Nango, S. Iwata, J. Standfuss, R. Neutze, in Annual Review of Biochemistry, Vol 88, Vol. 88 (Ed.: R. D. Kornberg), 2019, pp. 59–83.

[22] H. Qian, E. L. Elson, Proc. Natl. Acad. Sci. U.S.A. 2004, 101, 2828–2833.

[23] F. Garczarek, K. Gerwert, Nature 2006, 439, 109–112.

[24] T. Weinert, P. Skopintsev, D. James, F. Dworkowski, E. Panepucci, D. Kekilli, A. Furrer, S. Brunle, S. Mous, D. Ozerov, P. Nogly, M. T. Wang, J. Standfuss, Science 2019, 365, 61–65.

[25] J. K. Lanyi, Biochim. Biophys. Acta 1993, 1183, 241–261.

[26] M. Kataoka, H. Kamikubo, F. Tokunaga, L. S. Brown, Y. Yamazaki, A. Maeda, M. Sheves, R. Needleman, J. K. Lanyi, J. Mol. Biol. 1994, 243, 621–638.

[27] T. de Donder, P. V. Rysselberghe, Thermodynamic Theory of Affinity, Stanford University Press, California, 1936.

[28] S. R. de Groot, P. Mazur, Non-equilibrium thermodynamics, Dover, New York, 1962.

