## Supplementary Information for "Reaction cycle of operating pump protein studied with single-molecule spectroscopy"

**Materials.** Purified and lyophilized wt-bR and bR-D96N extracted from *Halobacterium salinarum* were purchased from Halotek biomaterials (Halotek, Germany). The sample was loaded into UV-treated 8-well chambers (Lab-Tek, ThermoFisher, USA) for single-molecule experiments. Single-molecule fluorescence experiments were carried out on a commercial Leica SP8x microscope (Leica Microsystems GmbH, Germany). Pulsed excitation light at 532 nm was provided by a Fianium Supercontinuum Laser operating at 40 MHz with an average power of ~ 6 μW at the back-focal-plane. Excitation and collection of fluorescence emission was carried out through a 100X oil-immersion objective. Fluorescence signal was passed through a band-pass filter (Semrock, USA) and was detected by photon avalanche photodiodes (PicoQuant, Germany) equipped with a picoHarp300 (PicoQuant, Germany) time-correlated single-photon-counting (TCSPC) system. Data processing and analysis was performed using in-house MATLAB code.

**Sample Preparation.** A 1 μM solution of membrane-bound bacteriorhodopsin (bR) was prepared in phosphate buffer (100 mM ionic strength, pH = 6). This solution was vortexed and sonicated for 5 minutes in cold water to disperse the purple membrane. The dispersed suspension was centrifuged for 5 minutes at 10K rpm. The supernatant was extracted, and the sonication/centrifuge procedure was repeated 3 times. The third-degree supernatant (~ 1 nM bR) was used for experiments. An 8-well chamber (LabTek, Australia) was UV treated for > 8 hrs prior to use. The bR solution was

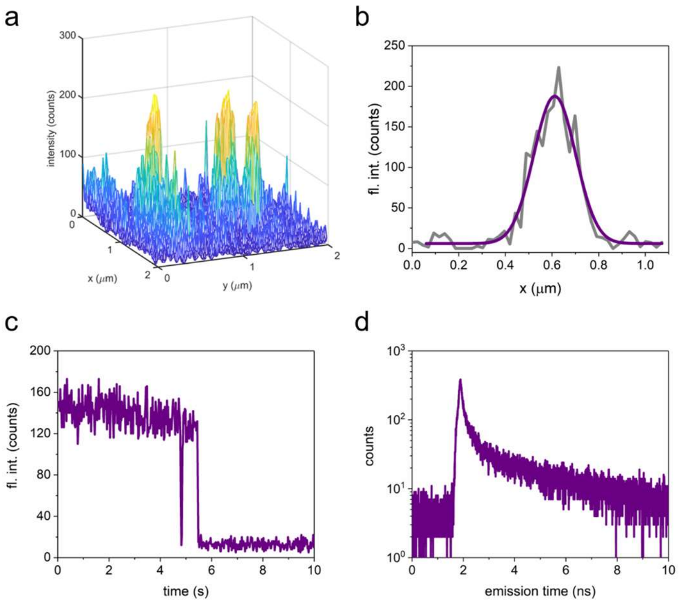


**Figure S1. Single-molecule imaging and photo-physics.** (**a**) Confocal image of single Bacteriorhodopsin proteins immobilized on a glass surface. (**b**) Diffraction-limited cross-section of fluorescence signal measured in confocal image. (**c**) Single-molecule fluorescence trace showing single-step photo-physics. (**d**) Fluorescence lifetime histogram from single wt-bR protein showing multicomponent relaxation.

then added to the chamber and kept at ambient light for 10-15 minutes. Membrane bound bR adsorb to the glass surface and are immobilized.

**SM-2D-FLCS.** Single-molecule dispersion and immobilization of bR was confirmed by confocal microscopy and fluorescence time traces (**Fig. S1a-c**). Emission from bR showed diffraction limited detection spots (**Fig. S1a, b**) and single-step photo-physics (**Fig. S1c**) expected for single molecules. Excitation at 532 nm overlaps with the absorption spectrum of the retinal chromophore (in ground state, K, L, and N intermediate states) embedded in the protein scaffold. The excitation laser both initiates the catalytic cycle and probes transient intermediates formed during the reaction cycle. Fluorescence relaxation from a single bR protein shows multi-component relaxation (**Fig. S1d**). The emission

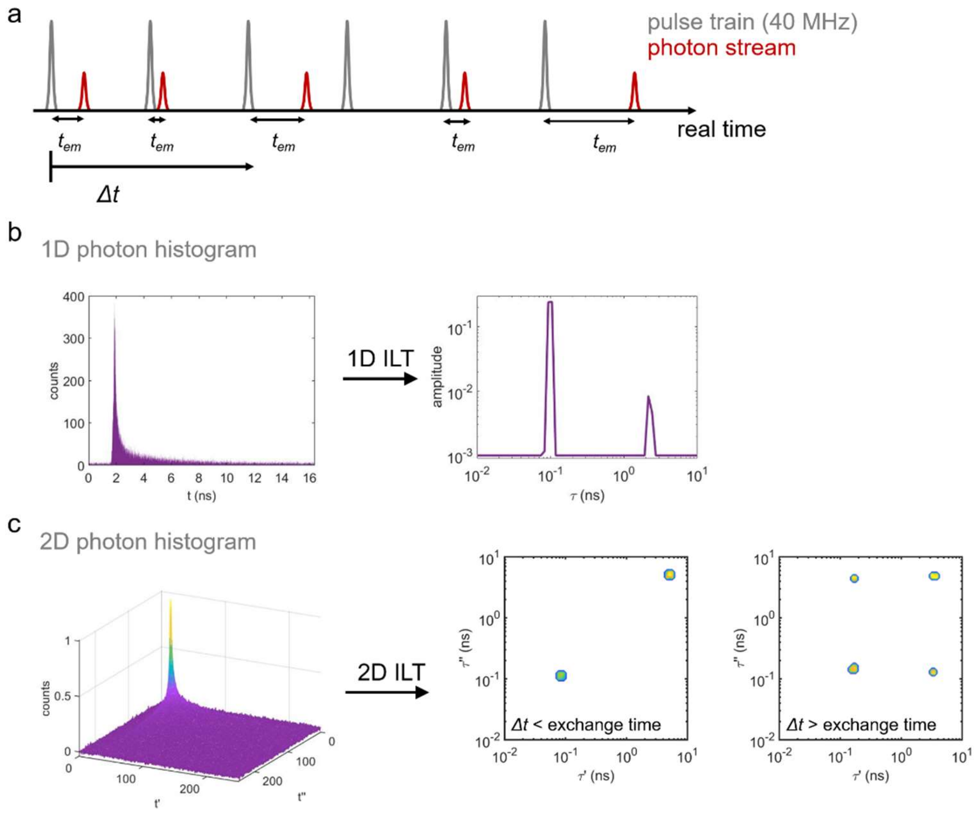


**Figure S2. Experimental procedure for measuring sm-2D-FLCS data.** (**a**) Excitation pulse train and photon stream measured with resolved photon emission time $t_{em}$ (standard TCSPC measurement in Time-Tagged-Time-Resolved mode). (**b**) 1D histogram of photon emission time and the fluorescence lifetime spectrum computed by a 1D ILT for a simple two state system. (**c**) 2D histogram of photon emission measured at a set waiting time $\Delta t$ and 2D-FLCS spectra computed by a 2D ILT. At early waiting times ($\Delta t$ < exchange time) peaks are only measured along the diagonal, indicating species that remained in their initial state during the waiting time. At later times ($\Delta t$ > exchange time), cross-peaks arise as species that were in an initial state exchange to the other state during the waiting time.

delay times extracted from a real-time photon stream (**Fig. S2a**) can be binned into a 1D histogram which are used to estimate a relaxation spectrum through an inverse Laplace transformation (**Fig. S2b**). For a two-state system where each state has a distinct fluorescence lifetime the 1D fluorescence lifetime spectrum will show a peak at the relaxation rate of each state. 2D-FLCS spectra are computed by generating a 2D photon histogram of photon pairs separated by a systematic waiting time $\Delta t$ (**Fig. S2a**). A 2D inverse Laplace transformation of the histogram gives a 2D-FLCS spectrum for a given $\Delta t$ (**Fig. S2c**). For a two-state system that undergoes chemical exchange at a well-defined timescale, the 2D-FLCS spectrum will contain diagonal peaks corresponding to species that did not exchange within $\Delta t$ and cross-peaks corresponding to species that did exchange within $\Delta t$ (**Fig. S2c**). Monitoring the kinetics of the cross-peaks provides direct access to the forward and reverse exchange times.

**State Occupancy.** The occupancies of the K, L, and N conformational states of bR was determined from 1D emission lifetime spectra. The relative amplitude of each state in the spectrum is proportional to the number of time steps in the experiment spent in each state. The occupancies of each state depend on temperature and change slightly over the temperature change used in this study (**Figure S4**).

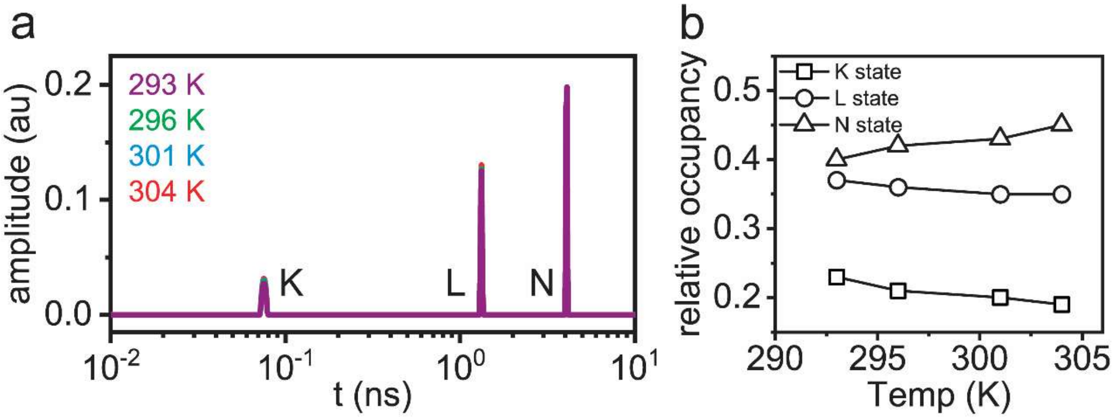


**Figure S3. Temperature dependence of bR state occupation.** (**a**) 1D fluorescence emission time spectrum and (**b**) relative state occupancies of bR as a function of temperature.

**Exchange Kinetics of D96N Mutant.** Exchange Kinetics of D96N Mutant. The L to N transition is observed to consist of an irreversible M1-M2 transition in wt-bR. The D96N mutant protein changes the aspartic acid residue at the 96 site, which is responsible for the proton uptake step, to a asparagine residue that hinders proton uptake (Fig. S5a). The D96N mutant has previously been observed to have a reversible switch step, with the M1-M2 and thus  effectively the L-N transition being significantly hindered.[1-2] Sm-2D-FLCS exchange experiments confirm the reversible nature of the L to N transition in the D96N mutant, without the forward transition timescale being significantly hindered.

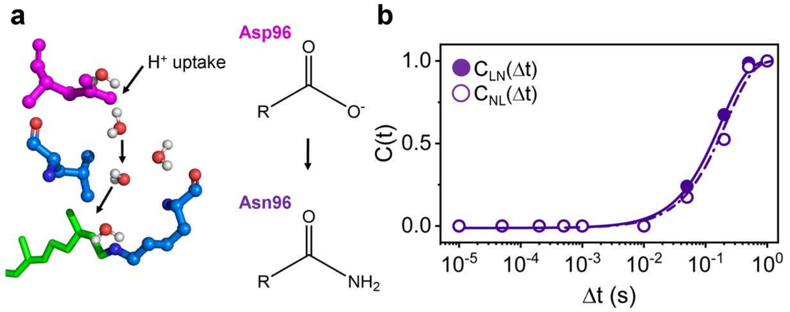


**Figure S4. Exchange dynamics of D96N mutant.** (**a**) Chemical structure of reprotonation site at Asp96 residue and the structure of the D96N mutant. (**b**) Chemical exchange from L to N (solid symbols) and N to L states (open symbols), showing near reversible exchange in the mutant protein.

**Inverse Laplace Transformations.** 1D and 2D inverse Laplace transformation (ILT) of the measured photon streams were performed through in-house MATLAB code described elsewhere.[3] From a photon stream collected from a single-molecule a 2D histogram is generated where each pixel denotes the number of coincidence events within the time trace where a photon with emission delay of $t_2$ is observed after $\Delta t$ from observation of a photon with delay time $t_1$. This represents the 2D emission delay correlation map, $X(t_1, t_2)$, which can be expressed as,

$$X(t_1, t_2) = \iint k_1(t_1, \tau_1) S(\tau_1, \tau_2) k_2(t_2, \tau_2) d\tau_1 d\tau_2 \qquad \text{(S1)}$$

where $S(\tau_1, \tau_2)$ is the 2D fluorescence correlation spectrum, $\tau$'s represents fluorescence lifetimes, and $k_1$ and $k_2$ are emission delay kernels. $S(\tau_1, \tau_2)$ is estimated by computing a 2D ILT of $X(t_1, t_2)$ at various $\Delta t$. This effectively decomposes the total correlation function into the autocorrelation function (diagonal elements) and cross-correlation function (off-diagonal elements) of the constituent components in the single-molecule fluorescence time trace. As

ILTs are numerically unstable and cannot be solved analytically, they instead must be estimated. Tahara and coworkers[4] used Maximum Entropy Method (MEM)[5] to estimate the 2D-FLCS spectrum, though this approach can be computationally costly. An alternative approach, based on recent advances in 2D NMR spectroscopy,[6-7] uses Tikhonov regularization[8] instead of MEM and exploits the kernel structure for efficient data compression via single-value decomposition, thus greatly reducing the computational costs without sacrificing the quality of fit of lifetime resolution.[3] We adapted this technique for the computation of sm-2D-FLCS spectra. For each $\Delta t$, the spectrum was computed using the photons within a time window for which the denoted $\Delta t$ was the upper limit (**see Photon Windows Section below**).

**Photon Windows.** sm-2D-FCLS spectra are computed using photons measured within a time window which is denoted by a single waiting time $\Delta t$. The time windows used for each waiting time are given below:

| *Time Window (s)* | *Denoted Δt (s)* |
|---|---|
| $1x10^{-6} – 1x10^{-5}$ | $1x10^{-5}$ |
| $1x10^{-5} – 5x10^{-5}$ | $5x10^{-5}$ |
| $5x10^{-5} – 2x10^{-4}$ | $2x10^{-4}$ |
| $2x10^{-4} – 5x10^{-4}$ | $5x10^{-4}$ |
| $5x10^{-4} – 1x10^{-3}$ | $1x10^{-3}$ |
| $1x10^{-3} – 1x10^{-2}$ | $1x10^{-2}$ |
| $1x10^{-2} – 5x10^{-2}$ | $5x10^{-2}$ |
| $5x10^{-2} – 2x10^{-1}$ | $2x10^{-1}$ |
| $2x10^{-1} – 5x10^{-1}$ | $5x10^{-1}$ |
| $5x10^{-1} – 1x10^{0}$ | $1x10^{0}$ |

**Influence of continuous laser illumination on the photocycle.** Bulk experiments using laser excitation has convincingly demonstrated that the photocycle mechanism of bR is not impacted by excitation[9]. Following photon absorption, a small amount of energy is dissipated as heat as the excited chromophore vibrationally relaxes. For low quantum-yield chromophores, however, the absence of fluorescence necessarily means that all the energy contained in the photon absorption must be thermally dissipated, leading to significant local heating. To demonstrate that this is not a concern in the bR experiments, we measured how the transition rates responded to small changes in external temperature (~18 ºC to 30ºC). As expected for an unperturbed system, the transition rates showed a strong dependence on changes to external temperature and nearly perfect Arrhenius behavior (**Figure 3. a,b**, main text). Such a clear trend would not be expected if the transition rates of the photocycle were strongly influenced from photon absorption during the probe process. This leads us to conclude that the dissipated energy from the probe does not interfere with the bacteriorhodopsin photocycle.

**Lapping effect in the photocycle.** We can rule out the lapping effect in the photocycle using sm-2D-FLCS experiments on bR-D96N mutant, which replaces the Asp96 residue responsible for proton uptake with Asn residue. This mutation is known to significantly hinder the L to N transition time (specifically, the M-N transition time), which contains the switch step responsible for the directional pumping action[10]. Hindering M to N with the D96N mutation causes this transition to become reversible. Our sm-2D-FLCS experiments show that the forward and reverse transition times for the L-N and N-L transitions are on the order of ~200 ms and 260 ms, respectively. The data is consistent with bulk experiments showing the reversibility of the switch step in the D96N mutant protein. Our results demonstrate that the reversibility is due to the destabilization of the N state hindering the L-N transition, while the reverse N-L transition is only slightly modulated. In the case of the D96N mutant the forward and reverse reaction times are comparable. Here, the only contribution to the N-L correlation function is through the direct reverse transition between the states and not through the lapping effect. The consistency between the timescale for the reverse transition rate from N-L in the wt-bR and the D96N mutant confirms that the lapping effect does not significantly contribute to the observed kinetics.

### References


[1] H. Luecke, B. Schobert, H. T. Richter, J. P. Cartailler, J. K. Lanyi, *Science* **1999**, *286*, 255-260.
[2] Y. Cao, G. Varo, M. Chang, B. F. Ni, R. Needleman, J. K. Lanyi, *Biochemistry* **1991**, *30*, 10972-10979.
[3] S. Talele, J. T. King, *Biophys. J.* **2021**, *120*, 4590-4599.
[4] K. Ishii, T. Tahara, *J. Phys. Chem. B* **2013**, *117*, 11414-11422.
[5] J. Skilling, R. K. Bryan, *Mon. Not. R. Astron. Soc.* **1984**, *211*, 111-124.
[6] L. Venkataramanan, Y. Q. Song, M. D. Hurlimann, *IEEE Trans. Signal Process.* **2002**, *50*, 1017-1026.
[7] Y. Q. Song, L. Venkataramanan, M. D. Hurlimann, M. Flaum, P. Frulla, C. Straley, *J. Magn. Reson.* **2002**, *154*, 261-268.
[8] A. N. Tikhonov, *Dokl. Akad. Nauk SSSR* **1963**, *151*, 1035-1038.
[9] J. K. Lanyi, *Annu. Rev. Physiol.* **2004**, *66*, 665-688.
[10] G. Varo, J. K. Lanyi, *Biochemistry* **1991**, *30*, 5008-5015.
